# Colonization of vineyards by non-*Saccharomyces* yeast species without evolution of copper and sulfite resistance

**DOI:** 10.1101/2025.09.22.677406

**Authors:** Justin C Fay, James H Miller, Sofia Dashko, Katie E Hyma, Ping Liu, Helena Volk, Haley Cohen, Nabil F Rahman, Keyang Tang, Emery R Longan, Xueying C Li, Lorena Butinar, Jure Piškur

**Author notes:** Present affiliations: H. Volk, Department of Agronomy, Biotechnical Faculty, University of Ljubljana, Ljubljana, Slovenia; X. Li, Ministry of Education Key Laboratory for Biodiversity Science and Ecological Engineering, College of Life Sciences, Beijing Normal University, Beijing, China, 100875; K. Tang, Department of Molecular and Cell Biology, University of Connecticut, Storrs, CT 06269; N. Rahman, Department of Pathology, University of Rochester Medical Center, Rochester, NY 14642. Deceased (Jure Piškur).

## Abstract

Humans have generated ecological and environmental disturbances, such as vineyards, across the globe. Disturbed environments create widespread and repeated selective pressures that can drive colonization and local adaptation in microbial species. We investigated the distribution of fermentative yeast species in vineyards compared to nearby arboreal habitats and measured their resistance to two commonly used vineyard antimicrobials, copper and sulfite. We analyzed 4,101 strains, representing 70 species, collected from grapevine- and oak-associated substrates at 17 vineyard and 20 non-vineyard sites in the USA and Slovenia. Species frequency varied with geography and substrate, but the majority of species commonly present in vineyards were also found in non-vineyard arboreal environments, representing a potential source for vineyard colonization and exploitation of sugar from grapes. Species varied in both copper and sulfite resistance, but only *Saccharomyces cerevisiae* showed elevated resistance in vineyard compared to non-vineyard samples. Our results indicate that *S. cerevisiae* has uniquely taken advantage of vineyard environments through adaptations that appear either unnecessary or inaccessible to other yeast species present in vineyards.

## Introduction

Humans have created a patchwork of disturbed environments across the globe (Haddad et al. 2015; Jacobson et al. 2019). From roadsides to forests, disturbed environments alter the abundance and distribution of native species and can result in the introduction and spread of exotic species (Lázaro-Lobo & Ervin 2019; Bowd et al. 2022). As a consequence, species have faced many new or altered selective pressures with the rise of agriculture and human population density (Turner 2010; Cheptou et al. 2017). Because human disturbances are often systematically applied across wide geographic areas, multiple species and populations can be subject to similar selective pressures. For example, the widespread application of herbicides and insecticides has resulted in repeated evolution of resistance both within and between diverse species (Bass et al. 2015; Wilson et al. 2025; Heap & Duke 2018).

Vineyards have been established around the globe for the production of grapes and wine. Similar to the expansion of other agricultural disturbances, vineyards arose subsequent to the domestication of grapes eleven thousand years ago (Dong et al. 2023), and now cover 7.1 Mha (17.5 million acres) of land globally (OIV 2024). Beyond the effects of land and crop management, vineyards also generate an energy rich but ephemeral source of sugar that can be scavenged by birds, insects and microbes.

Within budding yeasts of the Saccharomycetaceae family, Crabtree-positive yeasts ferment sugar even in the presence of oxygen and are well suited to take advantage of vineyard environments. The evolution of this fermentative lifestyle occurred over 100 million years ago coincident with the evolution of fruits and flowering plants (Pfeiffer & Morley 2014; Hagman et al. 2013). The advantage of fermentation over respiration can be attributed to a faster but less efficient use of sugar for energy production or to a niche construction strategy whereby sugar is metabolized to ethanol to inhibit competitors and then subsequently consumed (Merico et al. 2007). Grapes provide a concentrated source of fermentable sugar and domestic grapes ripen more synchronously and with higher sugar content than their wild relatives (Lu et al. 2024; Falginella et al. 2025). When ripe grapes are harvested and crushed, there is a diverse microbial community present at the start of fermentation (Bokulich et al. 2014; Gayevskiy & Goddard 2012). After fermentation, the wine is pressed and the pomace (seeds, skins) is disposed of or used to fertilize the vineyard, thereby completing a cycle of harvest, fermentation and vineyard re-inoculation (Sipiczki 2016). Thus, vineyards constitute a massively parallel anthropogenic disturbance open to yeast colonization and adaptation.

The Crabtree positive species, *Saccharomyces cerevisiae*, dominates wine fermentations and has repeatedly colonized and adapted to the vineyard environment. Whereas many species are fermentative and can turn grape juice to wine, *S. cerevisiae* quickly establishes itself as the most abundant species during wine fermentations (Fleet 2008; Csoma et al. 2010; Di Maro et al. 2007). *S. cerevisiae*’s ability to dominate wine fermentations depends on a variety of environmental factors and the mechanisms of dominance depend on the species it is competing with (Albergaria & Arneborg 2016; Williams et al. 2015). *Saccharomyces paradoxus*, the sister species of *S. cerevisiae*, shares its ability to dominate grape juice fermentations but is sensitive to copper and sulfite, which are commonly used in vineyards and winemaking.

Vineyard populations of *S. cerevisiae* have repeatedly evolved copper and sulfite resistance. Copper is frequently used in vineyards as a fungicide against plant pathogens (Borkow & Gabbay 2005), and sulfite is used in winemaking as an antioxidant and to prevent spoilage (Divol et al. 2012). Vineyard-associated strains of *S. cerevisiae* are copper resistant due to tandem amplification of the metallothionein *CUP1* (Fogel & Welch 1982; Zhao et al. 2014) and sulfite resistant due to chromosome rearrangements upstream of the sulfite efflux pump *SSU1* (Pérez-Ortín et al. 2002; Zimmer et al. 2014; García-Ríos et al. 2019). For both chemicals, resistance evolved multiple times through independent mutational events. Copper and sulfite resistance are thus considered vineyard adaptations as they are predominantly found in vineyard strains of *S. cerevisiae* whereas *S. paradoxus* and wild strains of *S. cerevisiae* are uniformly sensitive (Longan & Fay 2024; Dashko et al. 2016; Warringer et al. 2011).

Many non-*Saccharomyces* species are also present within vineyards and wine must (Drumonde-Neves et al. 2021; Barata et al. 2012; Jolly et al. 2014). However, colonization from non-vineyard environments and adaptation to vineyards has only been investigated in a few cases (Chandra et al. 2020). Genetic analysis has shown distinct vineyard populations of *Torulaspora delbrueckii* (Albertin et al. 2014; Silva et al. 2023), *Lachancea thermotolerans* (Hranilovic et al. 2017, 2018), and *Brettanomyces bruxellensis*, the latter of which also shows high sulfite resistance (Avramova et al. 2018). In contrast, population structure is primarily geography-associated for *Starmerella bacillaris* (Raymond Eder et al. 2019; Masneuf-Pomarede et al. 2015) and *Hanseniaspora uvarum* (Albertin et al. 2015). Identifying signatures of colonization and adaptation depends on comparisons with strains that were not isolated from vineyard or anthropized environments. For example, *S. bacillaris* exhibits geographic structure but has predominantly been found on grapes and in wineries (Raymond Eder et al. 2019). Because populations can exhibit fine-scale structure, isolation and characterization of strains from nearby environments is a key but frequently missing element in studying wine and vineyard-associated yeasts.

Yeasts can colonize vineyards from the surrounding environments or from other vineyards via human-associated migration. *S. cerevisiae* and *S. paradoxus* show notably different patterns of population structure indicative of colonization. Consistent with colonization from surrounding environments, *S. paradoxus* exhibits fine-scale population structure whereby vineyard isolates are most closely related to strains isolated from surrounding arboreal habitats (Hyma & Fay 2013; He et al. 2022). In contrast, *S. cerevisiae* strains isolated from vineyards are either members of a European-wine population that can be found in vineyards across multiple continents, or members of wild, indigenous populations (Gayevskiy et al. 2016; Hyma & Fay 2013; Loegler et al. 2024). Members of the European-wine population are only occasionally found outside vineyards and they are frequently human-associated. These and other patterns of population structure are often attributed to human-associated migration (Peña et al. 2025; Gayevskiy et al. 2016; Fay et al. 2019; Barbosa et al. 2016; Ludlow et al. 2016).

The sources of yeasts colonizing vineyards are largely unknown and may differ depending on the species (Griggs et al. 2021). Arboreal habitats represent one potential source (Cordero-Bueso et al. 2011; Chandra et al. 2020). Temperate forests harbor a high diversity of yeast species and largely overlap with regions of viticulture (Mozzachiodi et al. 2022). Most of the *Saccharomyces* species and the oldest lineages of *S. cerevisiae* have been isolated from arboreal sources, including bark, decaying leaf material and associated soil (Duan et al. 2018; Lee et al. 2022; Alsammar & Delneri 2020; Boynton & Greig 2014). The prevalence of *Saccharomyces* species in arboreal environments is high, with rates of isolation frequently exceeding 10% (Mozzachiodi et al. 2022). However, the ecology of *Saccharomyces* species in arboreal habitats is uncertain. *Saccharomyces* have been found in association with sap fluxes (Naumov et al. 1998) and traces of sugar have been detected in oak bark (Sampaio & Gonçalves 2008). Aphid honeydew provides a source of concentrated sugar (Völkl et al. 1999), particularly melezitose which is well fermented by *S. paradoxus* (Brysch-Herzberg & Seidel 2017). Accounting for sampling bias, it is possible that *S. cerevisiae* is an opportunistic generalist that is dispersed by insects and humans (Stefanini et al. 2012) and could come from any number of sources (Goddard & Greig 2015). Similarly, other yeast species, such as *Lachancea thermotolerance* and *Torulaspora delbrueckii*, are prevalent in vineyards but have also been frequently isolated from diverse habitats and sources (Spurley et al. 2022; Hranilovic et al. 2017; Silva et al. 2023; Albertin et al. 2014).

In this study, we examine the distribution of yeast species in vineyards compared to nearby arboreal environments. Species frequency provides insight into which species have successfully colonized and potentially adapted to vineyard environments. However, species frequency also depends on the substrate or source of isolation. To distinguish relative abundance related to source and geographic location, we compare tree and grapevine-associated samples within vineyards to tree-associated samples outside of vineyards. To test whether there has been parallel adaptation to vineyard environments, we measure copper and sulfite resistance of vineyard and non-vineyard strains.

Our study is based on a large collection of 4,101 yeast strains previously isolated from 17 vineyards and 20 non-vineyard locations in the USA and Slovenia (Figure 1). From this collection we identified 70 operational taxonomic units and those that vary by geographic location, environment and isolation source. We find vineyard yeast species are often present in nearby arboreal environments but only *S. cerevisiae* shows higher levels of copper and sulfite resistance in vineyard compared to non-vineyard samples. Our results provide insight into colonization of vineyard environments and local adaptation associated with domestication.

**Figure 1.**
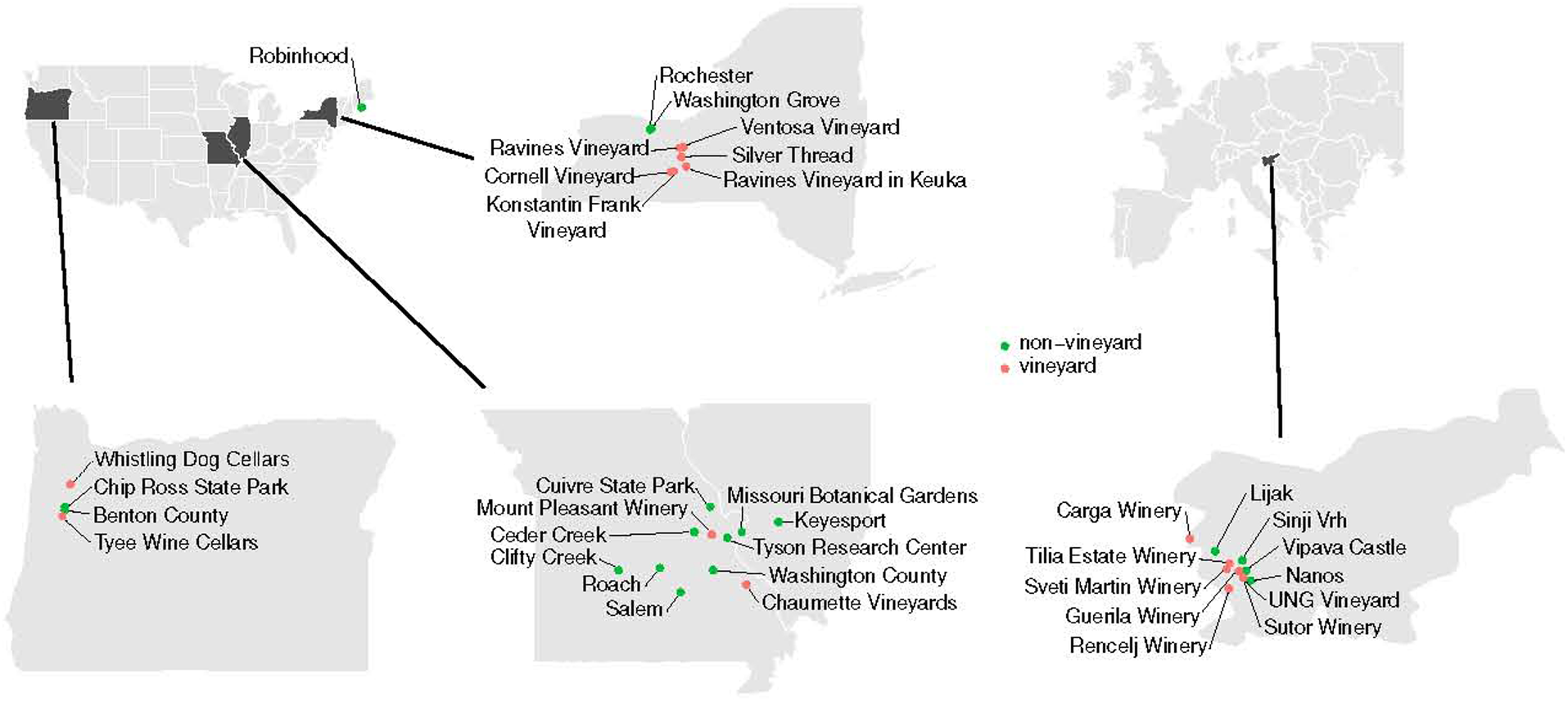
Sampling locations in the USA and Slovenia. Vineyard (red) and non-vineyard (green) sampling locations are shown for four magnified states in the USA and for Slovenia. Two non-vineyard locations in Hawaii are not shown.

## Materials and Methods

### Sample collections

Yeast strains were isolated from samples collected in the USA and Slovenia (Table S1). The first set consists of samples collected from four vineyards and four non-vineyard locations in Missouri and Oregon in 2008 and 2009 (Hyma & Fay 2013). The second set consists of samples collected from seven vineyards and four non-vineyard locations in Slovenia in 2013 and 2014 (Dashko et al. 2016). The third set of more heterogeneous samples consists of six vineyards and 12 non-vineyard locations in Missouri, Illinois, New York, Hawaii and Maine between 2012 and 2022. Samples were mainly of material associated with grapevines (soil, bark, berries) and various species of oak trees (soil, bark, twig) from both vineyard and non-vineyard locations. A small number of samples were of fruits, insects, pomace and wine must (Table S2).

### Yeast strains and isolation

Isolation of yeast strains was described previously and is briefly summarized here. Samples were enriched for fermentative yeasts at room temperature using a high sugar/ethanol enrichment medium (1% yeast extract, 2% peptone, 10% dextrose and 5% ethanol, adjusted to pH 5.3). A subset of samples was split and enriched using multiple methods. For Set 1 the samples were enriched using either a high sugar rich medium (above) or a low sugar minimal medium (6.7 g/L yeast nitrogen base with amino acids and ammonium sulfate, 1% dextrose and 8% ethanol). For Set 2, a subset of samples were split and enriched at both room temperature and 37 °C. For Set 3, a small number of samples were enriched by direct plating onto complete medium (CM) plates (6.7 g/L yeast nitrogen base with amino acids and ammonium sulfate, 2% dextrose, 2% agar) with different concentrations of copper sulfate or sodium sulfite. Enrichment cultures were streaked on rich medium (YPD) plates and a single colony was isolated and then stored as a stock at -80 °C. In some cases, multiple strains were isolated from the same enrichment when colonies showed visibly different morphologies.

A total of 4,101 strains were obtained from sampling. A set of 13 *S. cerevisiae* and nine *S. paradoxus* control strains representing known wild (North American Oak, Mediterranean Oak) and wine populations were added as controls (Table S3).

### Strain growth and phenotyping

Strains were grown from frozen glycerol stocks in 1 mL CM media arrayed in 96 deep-well plates and covered with sterile breathable rayon film for 3 days at room temperature. Grown cultures were resuspended by pipetting and 25 μL was transferred to PCR plates containing 5 μL 120 mM NaOH for cell lysis and 200 μL was transferred to standard 96-well plates for printing on CM plates.

Copper, sulfite, and temperature phenotyping were performed in three runs, where approximately 1,400 strains were phenotyped by colony size measurements in each run. For ease of handling, strains were kept in their 96-well orientations with specific wells reserved on four plates for control strains (Table S3). Printing to agar plates was handled robotically using a Singer Rotor HDA (Single Instruments, UK). The *S. cerevisiae* and *S. paradoxus* control strains were printed on each phenotyping plate in quadruplicate. Colonies were printed in a 1,536 arrangement by interleaving sixteen 96 colony plates. To control for colony size bias generated by plating location or neighboring colonies (Miller et al. 2022), two arrangements of each 1,536 plate were made. The first arrangement placed all control strains along the 31st row of the plate whereas the second rearranged the 96 colony plates so that all edge colonies from the first arrangement were interior colonies on the second and reshuffled the control colonies accordingly. Colonies were grown for two days at room temperature on CM plates between printings and were passaged at least twice at room temperature before printed on the plates used for phenotyping. Copper and temperature phenotyping plates consisted of CM agar with or without concentrated copper sulfate added to the flask after autoclaving, respectively. Sulfite phenotyping plates were created using a modification of a prior method (Park et al. 1999): CM agar was autoclaved with 75% the starting volume followed by the addition of sterilized sulfite buffer (150.3 mM L-tartaric acid, 150.3 mM sodium tartrate) to the desired final volume. As the solution cooled to handling temperature, sodium sulfite solution was added to obtain the desired final sulfite concentration. Copper and sulfite phenotyping plates were incubated for 36 hours at room temperature before photography in a Singer Phenobooth, whereas temperature phenotyping plates were incubated at their specified temperatures for 30 hours. Copper phenotyping plates were photographed using backlighting instead of the standard toplighting due to difficulties with accurate size quantitation of dark pigmented colonies which were common at intermediate and high copper concentrations. We also noticed that in many instances, *Schizosaccharomyces japonicus* was not pinned to phenotyping plates due to having crusty rather than sticky colonies. We therefore excluded this species from the phenotyping analyses.

Control strains were used to normalize the phenotype data across the three runs. Normalization was based on the average of ∼192 values from four technical plate replicates and 12 of the control strains that were present in quadruplicate on each plate. Normalization was separately performed for each stress and phenotype measurement. Following normalization, the four technical replicate values were averaged to obtain the final phenotype data.

Copper, sulfite and temperature resistance phenotypes were derived from colony sizes across stress treatments. There were 11 concentrations of copper, 16 concentrations of sulfite and 8 temperature treatments. Resistance to each stress was measured by the stress level at which growth was inhibited 50% (I50) as well as by the area under the curve (AUC) of colony size across stress levels. I50 was estimated by n-parameter logistic regression of the dose-response relationship using the nplr R package (Commo et al. 2025). To facilitate I50 fitting, colony sizes were adjusted to the largest colony size observed at higher stress levels. Colony size can increase with stress when competition for nutrients with adjacent colonies is eliminated due to their sensitivity to stress. AUC values were normalized relative to colony size in the absence of stress treatments. Differences in resistance between grapevine and non-vineyard oak samples were tested using a Wilcoxon signed-rank test with Holm-Bonferroni correction.

Additional phenotypes were measured for a subset of 589 strains from Set 1 (MO, OR) and 628 strains from Set 2 (Slovenia). Ethanol tolerance was measured on CM plates made with between zero and 18% ethanol with 1% increments after 36 hours of growth. Ethanol resistance was measured by the normalized and corrected area under the treatment-response curve. Growth on different carbon sources was measured using CM plates made with 2% ethanol, glycerol, fructose, galactose, maltose, maltotriose, melezitose, melibiose and sucrose. Growth on different carbon sources was measured by colony size relative to size on glucose plates. After removing strains of *Schizosaccharomyces japonicus* and strains not isolated from grapevines or oaks there were 1,053 strains.

### Species identification

Strains were identified to the species level by PCR amplification of the ITS2 region and sequencing. Overnight CM cultures were combined with 20 mM NaOH in PCR plates, covered with aluminum foil and frozen to -80 °C for at least 10 minutes, then incubated at 98 °C for 10 minutes in a PCR thermal cycler. After mixing by pipetting, 2 μL lysates were transferred to 18 μL PCR master mix containing 0.5 μM ITS2 binding primers with adapters compatible with the iNEXT barcoding system (Glenn et al. 2019): TCGTCGGCAGCGTCAGATGTGTATAAGAGACAG<u>GTGARTCATCRARTYTTTG</u> and GTCTCGTGGGCTCGGAGATGTGTATAAGAGACAG<u>CCTSCSCTTANTDATATGC</u> where the underlined regions are for ITS2 binding. PCR was performed in 96-well plates for 30 cycles of 30 second denature and anneal stages at 95 and 50 °C, respectively, followed by a 1 minute 72 °C extension. A 7 μL aliquot from the PCR reaction was cleaned of unbound primers by adding 0.5 μL exonuclease I (NEB M0293S) and 1 μL recombinant shrimp alkaline phosphatase (NEB M0371S) and incubated in the thermal cycler at 37 °C for 15 minutes followed by 80 °C for 15 minutes to deactivate the enzymes. The ITS2 amplicons were indexed using 0.5 uM iNEXT i5 and i7 barcoding primers (Glenn et al. 2019) in 25 μL PCR reactions containing 2.5 μL enzymatically-cleaned templates. Indexing was performed in 8 cycles of 95, 55 and 72 °C at 30 seconds each. Indexed libraries were combined for each plate (4 μL per reaction) and cleaned using a column-based purification kit (Qiagen 28104). All samples within a single run (1,536 samples) were combined and submitted as a single reaction for 2x300 paired-end sequencing on an Illumina MiSeq at Duke University’s sequencing core.

Sequencing reads were processed using programs from USEARCH v.11 (Edgar 2010; Edgar & Flyvbjerg 2015; Edgar 2016b). Reads containing PhiX sequences were removed with -filter_phix and then processed with Trimmomatic v0.39 (Bolger et al. 2014) with ILLUMINACLIP:adapters:2:30:10 SLIDINGWINDOW:4:20 LEADING:3 TRAILING:3 HEADCROP:22 MINLEN:150 where adapters sequences are TCGTCGGCAGCGTCAGATGTGTATAAGAGACAG and GTCTCGTGGGCTCGGAGATGTGTATAAGAGACAG with their reverse complements. Trimmed read pairs were merged with a maximum of five differences and 98 percent identity within the overlap region. Merged reads and surviving single reads with a minimum length of 150 bases and an expected error rate of 0.01 (Edgar & Flyvbjerg 2015) were denoised with unoise3 for reads with 8 replicates or more and alpha set at 7 (Edgar 2016b). The resulting sequences had chimeras removed by uchime3_denovo (Edgar 2016a) and clustered at 100% identity with cluster_smallmem into representative sequences of each sample (centroids). These centroids were cross listed against the samples to create an otutable and searched against a reference database to identify species. The best match was identified using ublast (Edgar 2010) with hits limited to at least 95% query coverage against either strand. We used the UNITE + INSDC reference database for fungi v. 8.3 (Abarenkov et al. 2025) with the following modifications. The ITS2 portion of each sequence was extracted using ITSx v.1.1.3 (Bengtsson-Palme et al. 2013a). All database entries that did not contain species level identities were removed. *S. cerevisiae* and *S. paradoxus* database entries that did not fully cover the ITS2 sequenced region were removed. Reference sequences from NCBI were added for samples that had less than 97% identity to the original database. Read counts and identity with reference hits associated with each species detected in a sample were compiled using custom scripts. For quality control, species identification was only used if it fulfilled the following criteria: at least 100 reads had species-level identities, at least 98% average identity to a referenced target sample-wide, 80% or more of the identified reads were associated with the same species (primary species). Applying these filters eliminated identification of 231 samples.

### Species labeling

Some species and strains were relabeled from their database matches (Table S4). Three species were relabeled using synonyms described in MycoBank (www.mycobank.org). Four species were relabeled Genus_species_group as we found closely related legitimate species that were indistinguishable in the ITS2 sequenced region using data from MycoBank and CBS (Vu et al. 2016). Using phylogenetic trees, 26 strains from ten species were relabeled Genus_unclassified if they did not closely match known species but branched between known members of a Genus. *Pichia* species were handled differently as there is substantial variation within legitimate species and they are often interspersed among each other (Wu et al. 2016). All centroids identified as *Pichia* species were clustered at 99.5% identity using cluster_smallmem from USEARCH and resolved into 19 distinguishable groups. These were aligned by PRANK v.170427 (Löytynoja 2014) to all Pichia references present in both the Utax reference database and a CBS database (Vu et al. 2016) where the ITS2 region was extracted using ITSx (Bengtsson-Palme et al. 2013b). Pichia clusters were relabeled using the species they most closely resembled on a parsimony tree. Using the tree, we labeled ten operational taxonomic units (OTUs) as *Pichia manshurica* and three as *Pichia membranifaciens*. Our rationale for multiple OTU labels was that they would be more conservative than lumping disparate groups when comparing copper and sulfite resistance in grapevine relative to other samples.

### Species validation

Species identification was validated using sequencing duplicates and comparisons to independent species identification. After quality control, there were 740 duplicates (strains with independent sequence identification) and 11 of these showed mismatches (1.5%). The mismatches involved five strains and species labels were removed. For independent validation, we used 17 control strains, 47 strains that were identified by Sanger sequencing of the ITS region, and 3,197 PCR/digests that identified *Saccharomyces* species and distinguished them from non-*Saccharomyces* species (Nardi et al. 2006; McCullough et al. 1998). Of these, 23 (0.71%) were not validated and species labels were removed. Excluding controls and unclassified strains, this left us with 3,812 strains representing 70 OTUs with the majority being identified to the species level (Table S5).

### Species frequency

To compare species frequency, we used a reduced set of strains that were consistently sampled from vineyard and non-vineyard locations. We removed strains from Set 3 as vineyard and non-vineyard samples were not collected at the same time and were not from the same regions (Table S1). For the remaining USA (Set 1) and Slovenia (Set 2) strains, we only used strains isolated from vineyard or non-vineyard oaks (soil, bark, twig) and vineyard grapevines (soil, bark, grape). We eliminated a small number of samples from 37 °C enrichments (Set 2). For USA samples (Set 1), strains collected in 2008 (samples 1-1084) were obtained from two rounds of high (H) or low (L) sugar enrichment media: HH, HL, LH, LL. In the first round, samples were immediately split into high or low sugar enrichments and so we consider strains from the HH and HL enrichments to be independent of the LH and LL enrichments. We therefore only used strains from the HH and LH enrichments. Because the Slovenia samples (Set 3) were only obtained from high sugar enrichments, this limited comparison of USA and Slovenia samples. However, within the USA samples, where samples were put through both the HH and LH enrichments, only three of the 40 species showed significantly different rates of isolation between the HH and LH enrichments (Fisher’s Exact test, p < 0.01, Table S6). The reduced strain set consisted of 2,364 strains from non-vineyard oaks (n = 858), vineyard oaks (n = 891) and grapevines (n = 615) in Missouri (n = 1,188), Oregon (n = 524) and Slovenia (n = 652)(Table S7).

Species frequency was evaluated using logistic regression. We used Firth’s bias reduction method implemented with the logistf (v1.26.0) R package (Heinze et al. 2025). The method uses a penalized maximum likelihood to avoid problems of separation and infinite parameter values that occur when there are zero species counts in a particular category or interaction (Heinze & Schemper 2002). Species frequency was fit to a model of sample source (grapevine, vineyard oak, non-vineyard oak) and pairwise comparisons were tested using the emmeans (v1.10.0) R package (Lenth et al. 2025) with Holm-Bonferroni adjustment for multiple comparisons (Table S8). Species frequency was also tested for association with geographic region (MO, OR, Slovenia) and for an interaction between region and sample source with Holm-Bonferroni adjustment for multiple comparisons (Table S8).

## Results

### Samples and species diversity

A total of 4,101 strains were obtained from 37 sampling locations in the USA and Slovenia (Table S1). Most of the strains were isolated from oak-associated substrates (bark, soil, twig) within vineyard (36%) and non-vineyard locations (38%), and from grapevine-associated substrates (bark, soil, grape) within vineyards (21%)(Table S2). We sequenced the ITS2 region and resolved 3,812 strains to 70 operational taxonomic units (OTUs, Table S5). The majority of OTUs matched 55 known species. However, four OTUs were only distinguished to the genus level due to closely related species and 11 OTUs related to the *Pichia manshurica/membranifaciens* species complex formed distinguishable groups based on their ITS2 region (Figure S1).

Most strains (93%) were from 17 common (>0.5%) species (Table 1). Five of these species were predominantly found in vineyard locations (>90%), and ten of the species were predominantly found in a single country. To compare species frequency between grapevines and oaks within and outside of vineyards, we used a reduced set of strains that were consistently sampled across time periods, locations and enrichments (n = 2,364, see Methods).

**Table 1.** Number of strains isolated by location.

| Species | Group | Non-vineyard | Vineyard | USA | Slovenia |
| --- | --- | --- | --- | --- | --- |
| *Saccharomyces paradoxus* |  | 398 | 643 | 766 | 275 |
| *Saccharomyces cerevisiae* |  | 297 | 310 | 439 | 168 |
| *Lachancea thermotolerans* |  | 160 | 338 | 452 | 46 |
| *Lachancea fermentati* | USA | 193 | 142 | 335 | 0 |
| *Schizosaccharomyces japonicus* | USA | 163 | 140 | 303 | 0 |
| *Torulaspora delbrueckii* |  | 76 | 75 | 112 | 39 |
| *Pichia kudriavzevii* | Vineyard | 2 | 114 | 94 | 22 |
| *Starmerella bacillaris* | Vineyard-USA | 0 | 90 | 88 | 2 |
| *Hanseniaspora vineae* | Vineyard-USA | 8 | 75 | 83 | 0 |
| *Kazachstania servazzii* |  | 26 | 39 | 15 | 50 |
| *Pichia occidentalis* | Vineyard-USA | 0 | 50 | 50 | 0 |
| *Hanseniaspora uvarum* | Slovenia | 5 | 40 | 0 | 45 |
| *Kluyveromyces lactis* | USA | 20 | 20 | 39 | 1 |
| *Zygosaccharomyces bailii* | USA | 9 | 31 | 36 | 4 |
| *Kazachstania martiniae* | USA | 19 | 18 | 37 | 0 |
| *Wickerhamomyces anomalus* | Vineyard | 1 | 22 | 19 | 4 |
| *Saccharomyces kudriavzevii* | Slovenia | 13 | 8 | 0 | 21 |
| Rare species |  | 93 | 174 | 188 | 79 |
| Total |  | 1483 | 2329 | 3056 | 756 |

Groups indicate species with over 90% of strains from a vineyard or location in the USA or Slovenia.
Rare species is the sum of all species with fewer than 20 strains.

### Species frequency by location and sample source

We tested whether species distributions varied by sampling site (vineyard vs non-vineyard) or source (oak vs grapevine) for 16 species with at least 20 strains in the reduced set. To examine isolate sources while controlling for site, we compared species frequency from grapevines to vineyard oaks and found eight species were more abundant on grapevines and four species were more abundant on oaks (Figure 2A, Holm-Bonferroni p < 0.05, Table S7). Six of the eight species enriched on grapevines were predominantly grapevine-associated as they were not present or rare (< 1%) on oaks regardless of their location. The other two grapevine enriched species were *Lachancea thermotolerans* and *Lachancea fermentati* and they were both common on oaks. Thus, oaks serve as a potential reservoir for grapevine colonization for 10 of the 16 species.

**Figure 2.**
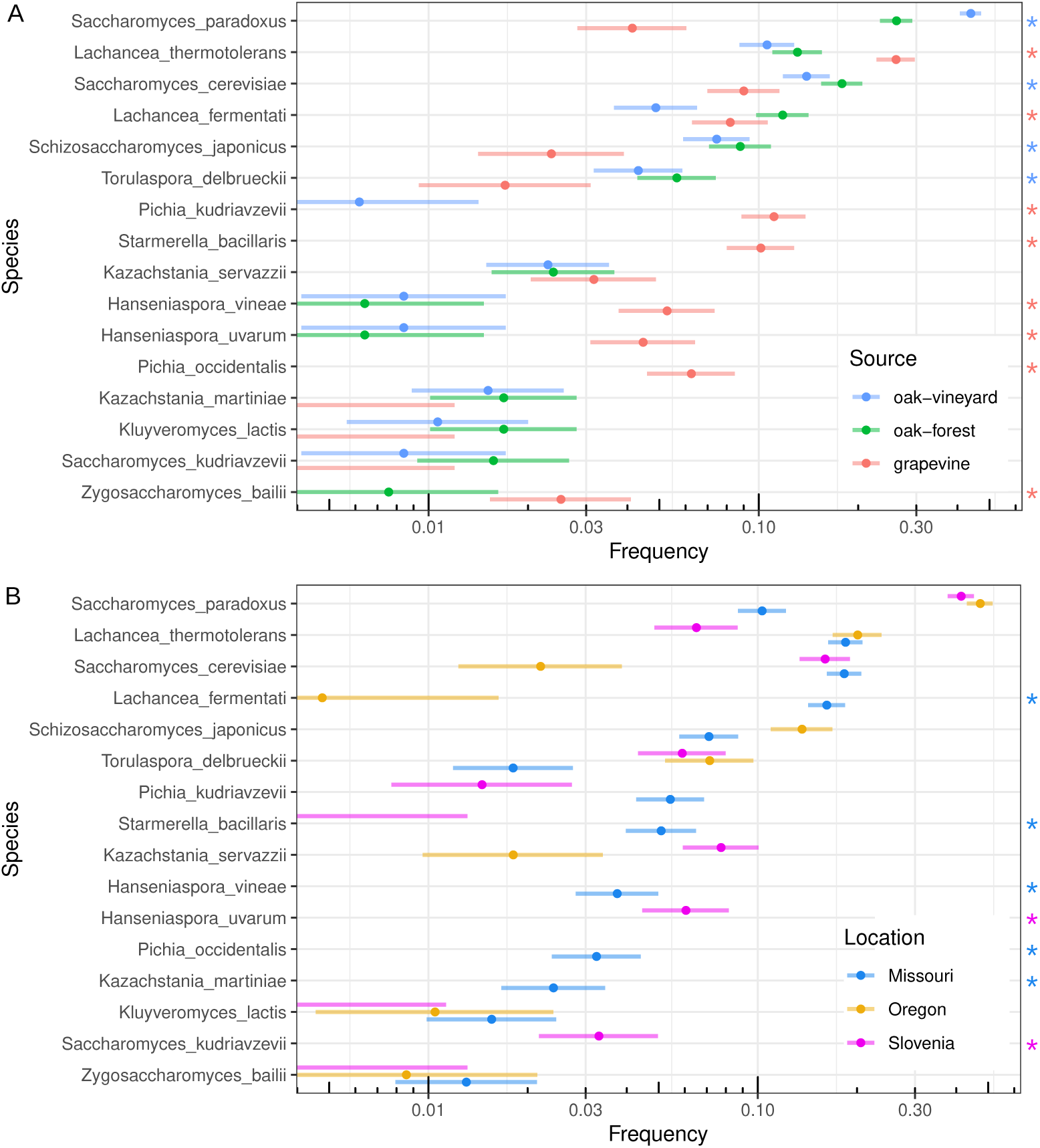
Species frequency by source and location. A. Species frequency for grapevines and oaks from both vineyard and non-vineyard locations. Asterisk (right) indicates species with higher frequency on grapevine (red) versus vineyard oaks (blue). B. Species frequency for Missouri, Oregon and Slovenia sampling locations. Asterisk (right) indicates species only found above 1% frequency in a single location. For both panels, bars show 95% confidence intervals and species are ordered by overall frequency.

To compare vineyard to non-vineyard sites while controlling for source we compared species frequency from vineyard oaks to non-vineyard oaks. Two species differed: *S. paradoxus* and *L. fermentati* (Figure 2A). However, neither of these species are predominantly vineyard-associated. *L. fermentati* was more abundant on non-vineyard oaks and *S. paradoxus* was the most abundant species on non-vineyard oaks and has been isolated from tree-associated samples across the globe (Boynton & Greig 2014; He et al. 2022).

Species frequency also differed across broad geographic locations. We compared species frequency in Missouri, Oregon and Slovenia and found all the species showed significant differences except *Zygosaccharomyces bailii* (Figure 2B, Holm-Bonferroni p < 0.05, Table S7). Furthermore, only four species were common (> 1%) in all three locations: *S. cerevisiae*, *S. paradoxus*, *L. thermotolerans* and *T. delbrueckii*, and seven species were only common in one of the three locations, including four of the predominantly grapevine-associated species.

Given the large differences in frequency across regions, we also tested for an interaction between geographic region and sampling site/source (grapevine, vineyard oak, non-vineyard oak). Four species showed significant interactions (Figure S2, Holm-Bonferroni p < 0.05, Table S7). *S. cerevisiae* showed higher frequency on grapevines compared to vineyard oaks in Slovenia but lower frequency in Missouri. *S. paradoxus* showed higher frequency in vineyard compared to non-vineyard oaks in Oregon and Slovenia but lower frequency in Missouri. *T. delbrueckii* and *L. thermotolerans* showed more complex patterns.

### Phenotypic divergence within vineyards

We measured copper and sulfite resistance to determine whether vineyard strains are more resistant than non-vineyard strains for each common species. Phenotypes of control strains (n = 22) showed that we could distinguish *S. cerevisiae* wine strains from non-vineyard oak strains of *S. cerevisiae* and *S. paradoxus* by their higher levels of copper and sulfite resistance (Figure 3A and 3B). Among the eight species with sufficient (n > 2) samples from both sources, *S. cerevisiae* was the only species with significantly higher copper or sulfite resistance in grapevine compared to non-vineyard oak samples (Figure 3C and 3D, Table S9). *S. cerevisiae* strains isolated from vineyard oaks showed intermediate resistance and spanned the range of copper and sulfite resistance levels, consistent with a mixture of strains from vineyard and wild populations (Dashko et al. 2016; Hyma & Fay 2013). Two species, *L. fermentati* and *L. thermotolerans*, actually showed higher copper resistance in non-vineyard compared to grapevine samples.

**Figure 3.**
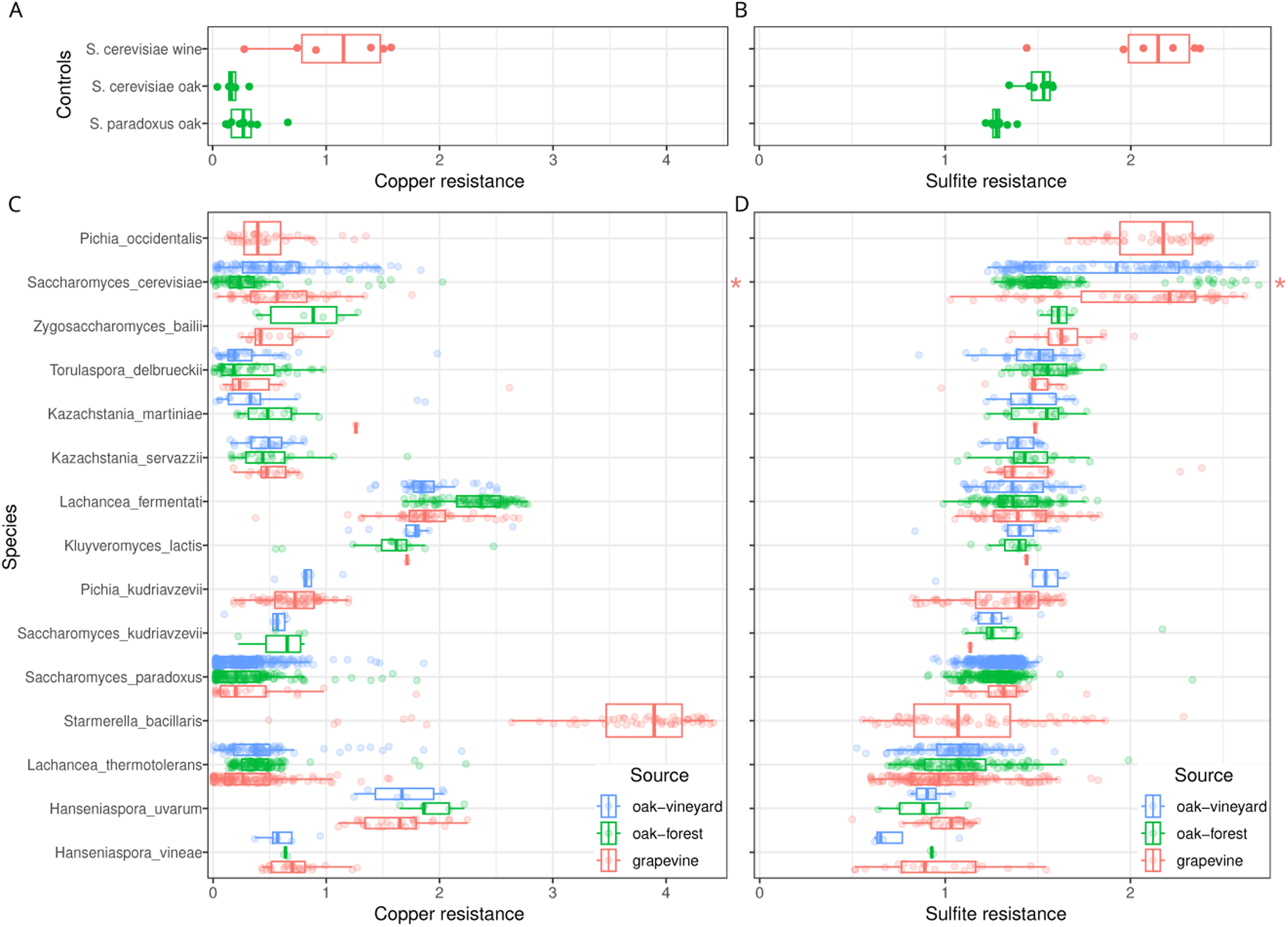
Variation in copper and sulfite resistance. Copper (A, C) and sulfite (B, D) resistance measured by the area under a curve formed from colony size in response to treatment. Strains (points) are colored according to whether they came from grapevine, oak-forest or oak-vineyard samples (source) and boxplots show median and interquartile range for phenotyping controls (A, B) and each species (C, D). Species were ordered with respect to sulfite resistance. Asterisk (right side of C and D) indicates species with significantly higher resistance of strains from grapevine compared to oak-forest samples.

Species varied widely in both copper and sulfite resistance and some were more resistant than *S. cerevisiae* wine strains. Four species (*L. fermentati*, *K. lactis*, *S. bacillaris* and *H. uvarum*) showed notably high resistance to copper, and the most resistant of these (*S. bacillaris*) was only found in vineyards. A single species, *P. occidentalis*, approached levels of sulfite resistance present in *S. cerevisiae* wine strains, and was also only found in grapevine samples.

For a subset of strains (n = 1,217), we also measured ethanol and temperature tolerance as well as growth on nine different carbon sources. We chose these traits as plausibly associated with vineyard adaptation. Wine must attains high levels of ethanol and vineyard soils can reach high temperatures due to direct sunlight exposure (Costa et al. 2023). Grapes contain high levels of fructose and glucose and yeast species vary in the breadth of their sugar utilization, with those in the *Wickerhamiella*/*Starmerella* clade evolving fructophily (Gonçalves et al. 2018; Opulente et al. 2024).

Species varied widely in their growth on different carbon sources and in their ethanol and temperature tolerance (Figure S3). However, only two species other than *S. cerevisiae* exhibited significant differences between grapevine strains and non-vineyard oak strains (Holm-Bonferroni p < 0.05, Table S9). *L. thermotolerans* grapevine strains showed reduced growth on galactose and at high concentrations of ethanol. *Kazachstania servazzii* grapevine strains were more ethanol tolerant than non-vineyard oak strains. *S. cerevisiae* showed multiple phenotype differences. *S. cerevisiae* grapevine strains showed higher growth on maltose and reduced growth on melezitose, galactose, ethanol, glycerol and at high temperature compared to non-vineyard oak strains.

## Discussion

The widespread cultivation of grapes for wine production generates a rich but transient source of sugar in vineyards. In this study, we surveyed the distribution of fermentative yeast species within and outside of vineyards and compared their levels of resistance to copper and sulfite. Many yeast species present in vineyards were also abundant in nearby arboreal environments but did not exhibit elevated resistance in vineyards. However, we also identified species that were predominantly found within vineyards, some of which displayed copper or sulfite resistance comparable to that of *S. cerevisiae* wine strains. Below we discuss both ecological and evolutionary factors relevant to species’ colonization of vineyards and signatures of vineyard adaptation.

Our survey of yeast species is largely consistent with prior studies of species isolated from vineyard and non-vineyard environments. Out of 16 species abundant in our samples, all but two were previously isolated from grapes or wine must (Drumonde-Neves et al. 2021): *Kazachstania martiniae* and *S. japonicus*. Both of these species were predominantly obtained from oak samples and only found in the USA. Similarly, 12 out of 16 species were also isolated in a large survey of soil, plants, trees and other substrates in the USA (Spurley et al. 2022). The four species not previously found consist of two grapevine-associated species, *S. bacillaris* and *Z. bailii*, and two oak-associated species, *S. japonicus* and *Saccharomyces kudriavzevii*, the later of which was only found in Slovenia.

Our sampling scheme enabled us to identify yeast species that varied by source (grapevine vs oak-associated) and by location (vineyard vs non-vineyard) without confounding between these factors. We found that most species varied by sample source and/or location. To summarize large differences we delineated species enriched in grapevine compared to vineyard oak samples and species enriched in the USA compared to Slovenia using an odds ratio of three or more (Figure 4, Table S7). Out of 16 species, six were grapevine-enriched, five were oak-enriched and eleven were enriched in either USA or Slovenian samples. Thus, at a local scale of vineyard versus non-vineyard, sample source is the better predictor of species frequency, but over broader regions (Missouri, Oregon, Slovenia) location plays a dominant role. One notable feature of the geographic differences is that all of the USA enriched species except *S. japonicus* were almost entirely found in Missouri and not Oregon, suggesting that climate or other ecological variables are more important than geographic distance.

**Figure 4.**
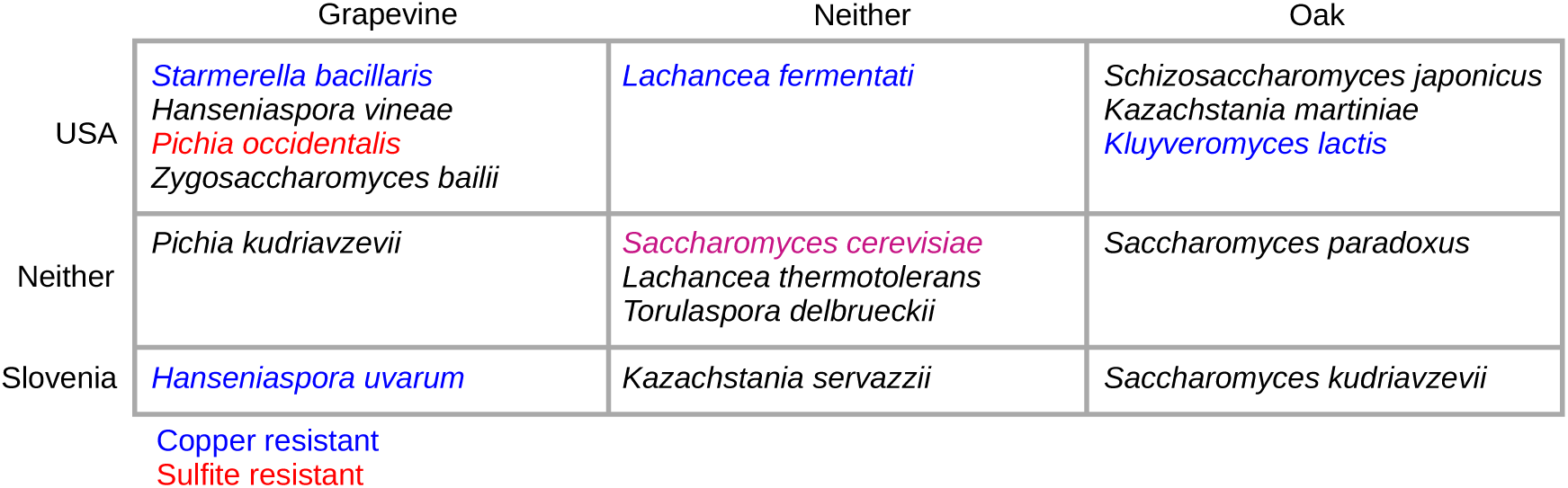
Summary of species distributions. Species were delineated as grapevine or oak enriched, USA or Slovenia enriched, or not enriched (neither) using an odds ratio of three. Species with average levels of copper (blue) or sulfite (red) resistance at or above *S. cerevisiae* wine strains (Figure 3) are color coded. Enrichments were all significant except for the four least abundant species (Table S7).

Our survey shows that arboreal habitats provide a possible source population for colonization of vineyards. Out of 16 commonly recovered species, only three were not found outside vineyards: *Pichia kudriavzevii*, *Pichia occidentalis* and *S. bacillaris* (Table S7). While other species were enriched in grapevine samples, these species may be transiently abundant in vineyards depending on the time of year. Thus, continuous migration from arboreal populations into vineyards provides one explanation for species that show no evidence for local adaptation to copper and sulfite (Holt & Gaines 1992).

Fitness provides another explanation for species that have not evolved copper or sulfite resistance. The average level of copper or sulfite resistance of five species is as high as that present in *S. cerevisiae* wine strains (Figure 3). In these species there may be little or no benefit of evolving higher levels of resistance in vineyard populations. Other aspects of ecology may also limit fitness benefits. For example, *L. thermotolerans* can be eliminated from wine fermentations by antimicrobial peptides generated by *S. cerevisiae* (Kemsawasd et al. 2015; Branco et al. 2017), which could reduce or eliminate the benefits of sulfite resistance during wine fermentation.

Costs of resistance and genetic constraints on adaptation could also influence the evolution of copper and sulfite resistance (Longan & Fay 2024). *S. cerevisiae* wine strains are copper resistant due to *CUP1* duplication but *CUP1* is only present in the *Saccharomyces* species and may not serve as a mechanism of resistance in other species. Additionally, a laboratory derived *CUP1* duplication in *S. paradoxus* was found to be quite costly in the absence of copper (Longan & Fay 2024).

There are a variety of ways in which adaptation to vineyards would not be detected in our study. First, we relied on comparisons to oak-associated isolates, which were not found for some species. Second, we focused on copper and sulfite resistance whereas other traits may play more important roles in vineyard adaptation. Finally, migration and gene flow can obscure phenotypic signals of local adaptation unless population genetic structure present within a species is simultaneously examined. This likely contributes to the wide range of copper and sulfite resistances observed in *S. cerevisiae* strains isolated from grapevines (Figure 3). Population genetic analysis of vineyard and non-vineyard populations would help determine whether vineyard populations have become genetically differentiated due to local adaptation, and would also provide better insight into where vineyard strains came from.

In a broader context, our results support the view that domestication is restricted to certain species (Diamond 2002). Although the definition of domestication is debated (Lord et al. 2025), it generally entails mutual benefits between humans and the domesticated organism (Purugganan 2022). In this light, copper and sulfite resistance can be regarded as domestication traits: yeast gain access to high sugar environments, while humans benefit from the suppression of undesirable microbes during wine production. Our finding that the evolution of copper and sulfite resistance is rare in yeasts other than *S. cerevisiae* is thus particularly striking when compared to domesticated plants where similar genes underlie convergence evolution of domestication phenotypes across multiple species (Purugganan 2019).

## Acknowledgements

This work was supported by the National Institutes of Health (GM080669 to JCF), the European Regional Development Fund (3330-13-500031 to JP) and the Slovenian Research Agency (ARRS J4-4300 and BI-US/13-14-028 to JP and JCF). Samples were collected with the assistance of Katarina Koruza, Timothy Lai, Elizabeth Engle and Devjanee Swain Lenz.

## Data availability

Raw data, scripts and other files used to generate results are available at OSF (https://doi.org/10.17605/OSF.IO/ZDMRB).

---

**Figure S1.**
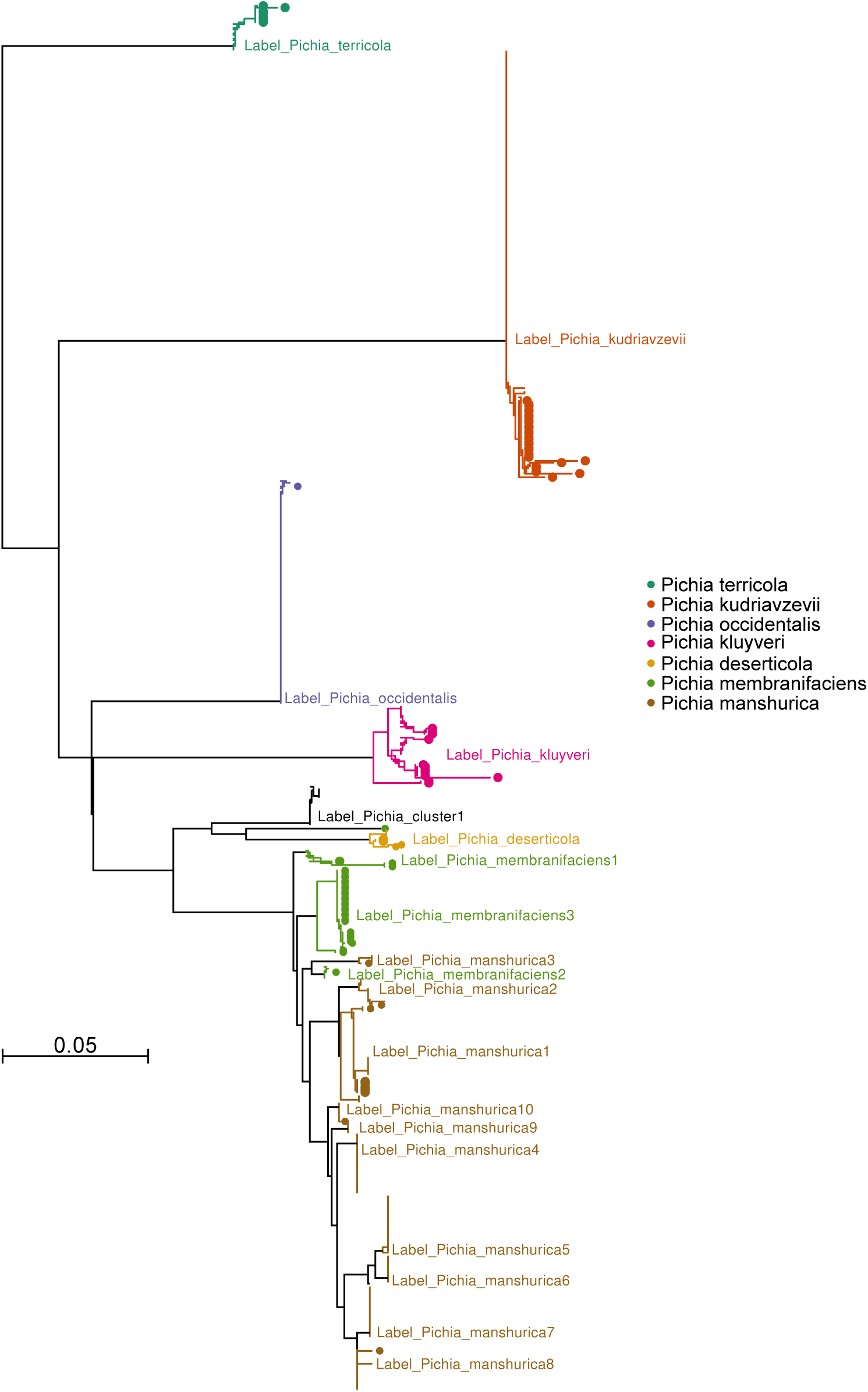
Phylogenetic relationship of *Pichia* species and sequenced strains. Phylogeny shows known species (circles) and strains (no label) based on ITS2 sequences. Strain branches are colored according to the species or group labels.

**Figure S2.**
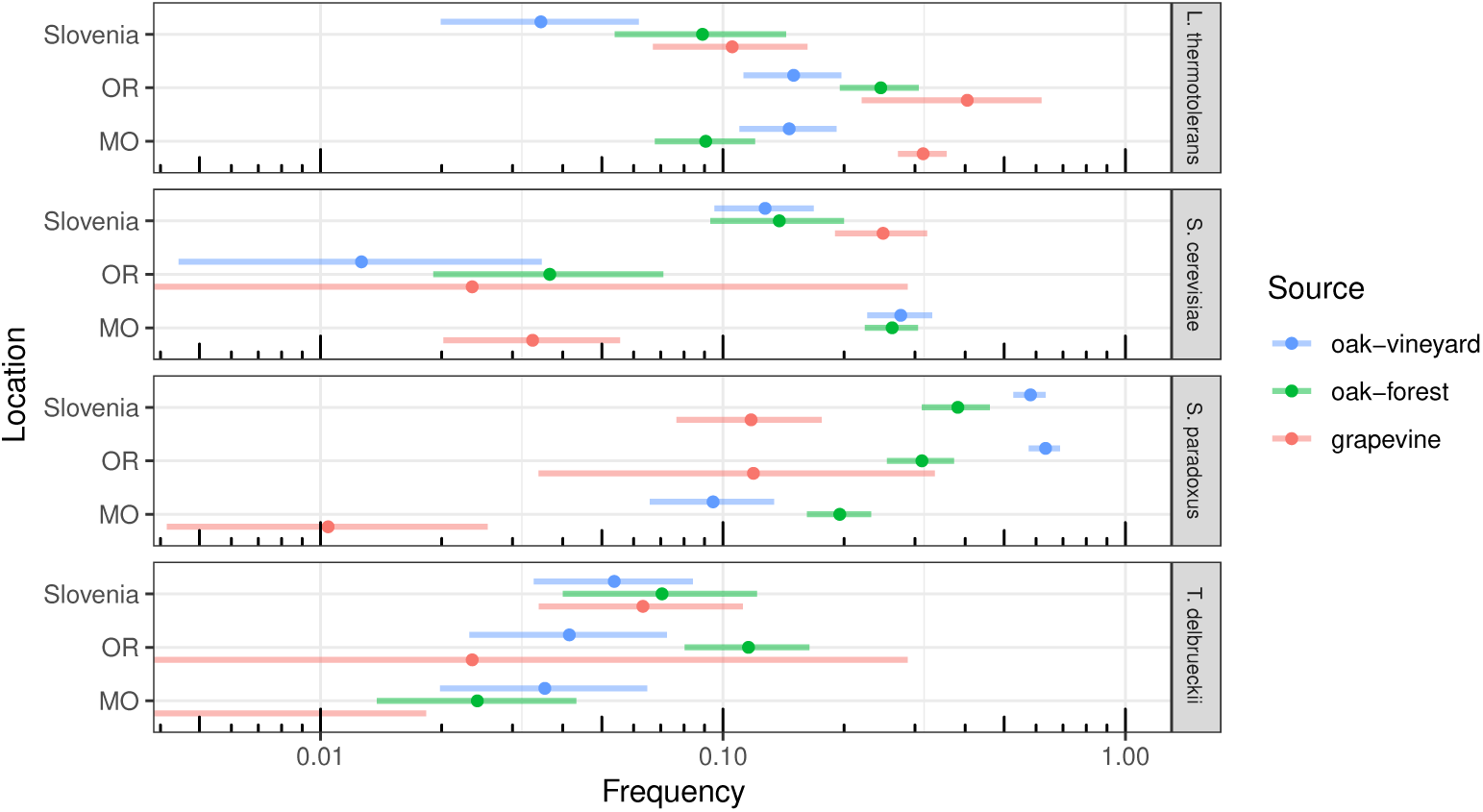
Species frequency across regions and sample sources. Each panel (species) shows species frequency in Missouri (MO), Oregon (OR) and Slovenia for samples of grapevines and vineyard and non-vineyard oaks. Bars show 95% confidence intervals.

**Figure S3.**
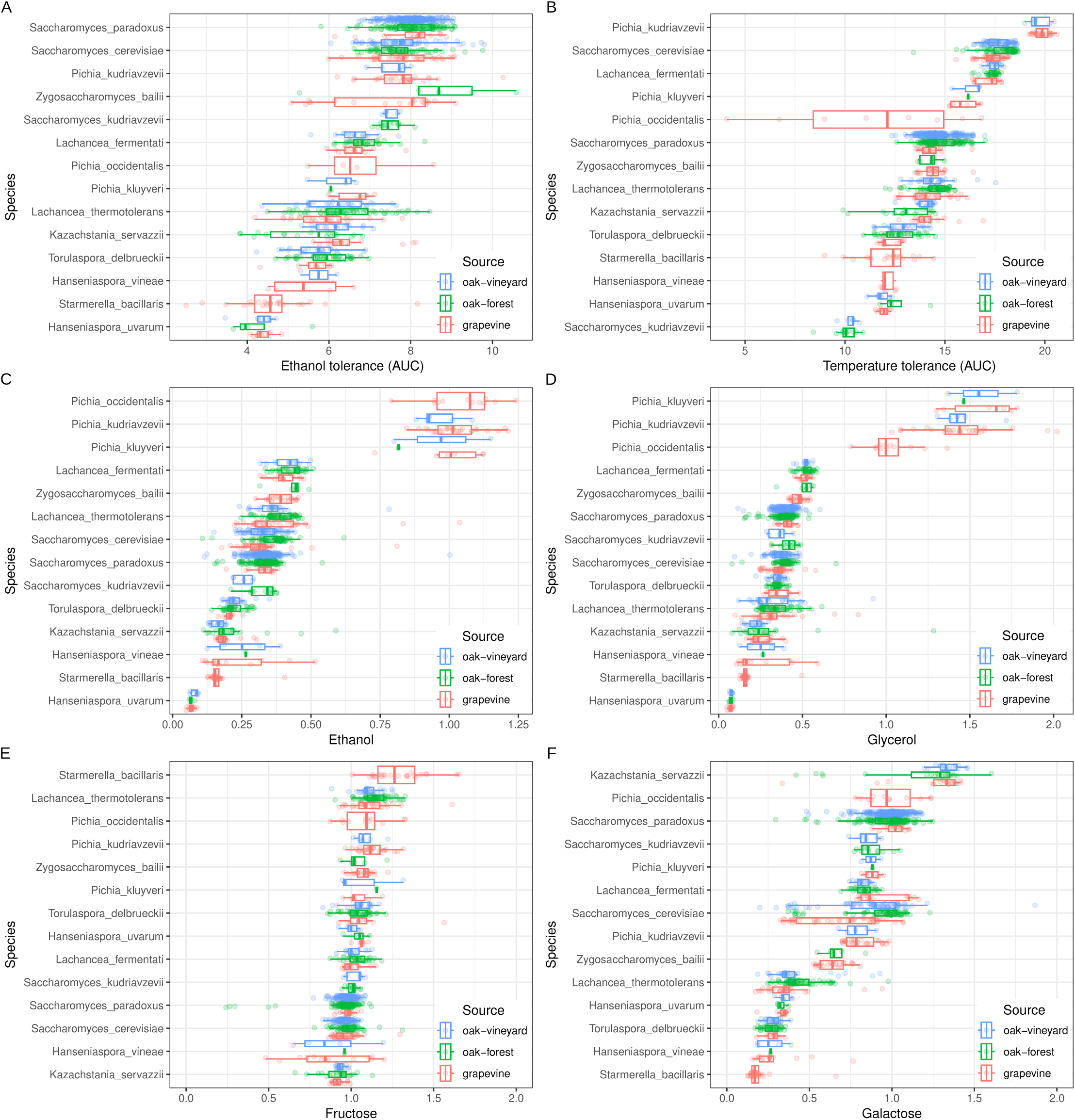
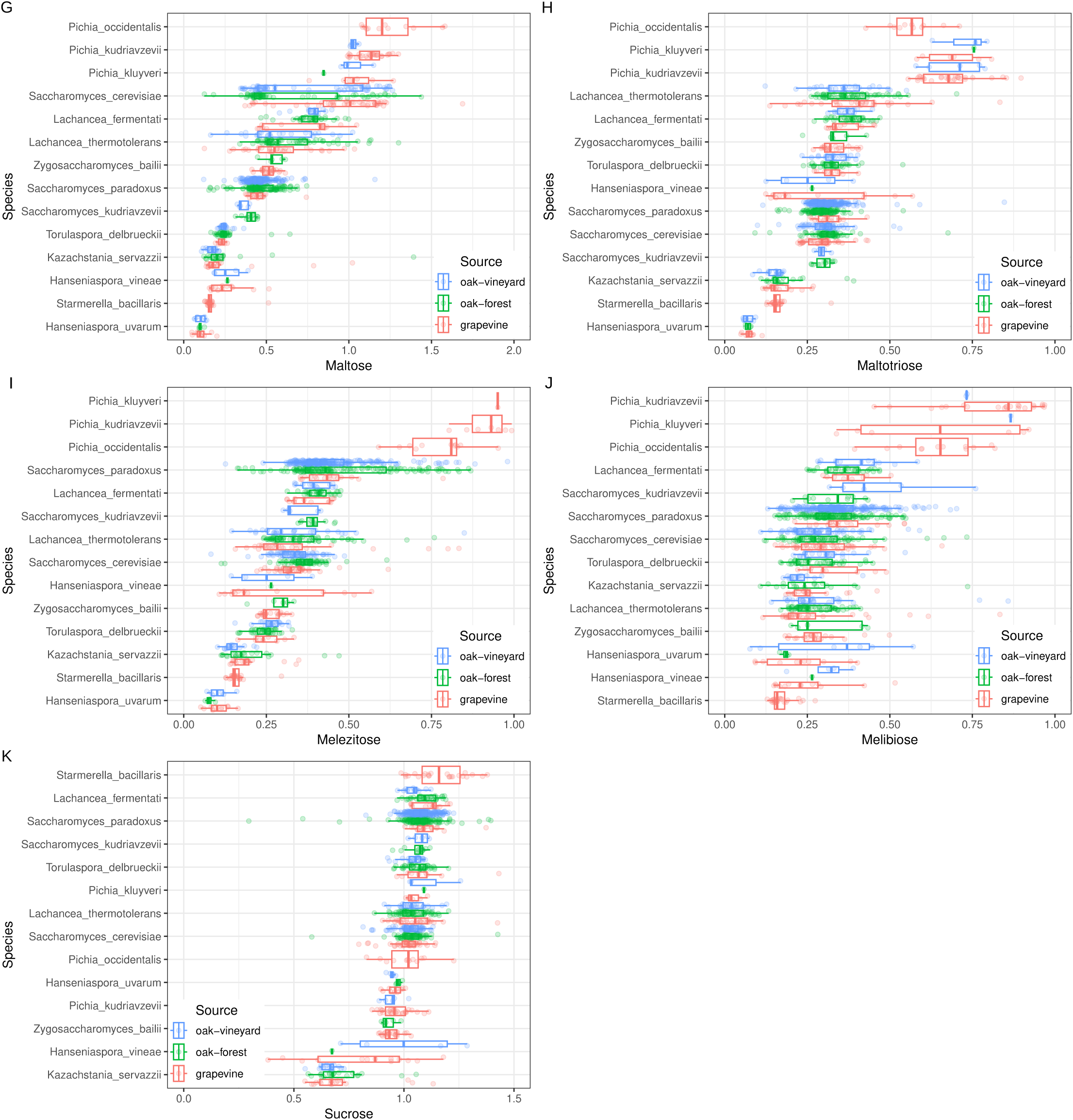
Phenotypic variation within and between species. Phenotypic variation is shown for 14 species with ten or more strains in the subset used for extended phenotyping. Ethanol (A) and temperature (B) tolerance was measured by area under the curve (AUC) normalized to no ethanol and 23 °C, respectively. On this scale AUC is in units of percent ethanol or degrees above 23. Growth on ethanol (C), glycerol (D), fructose (E), galactose (F), maltose (G), maltotriose (H), melezitose (I), melibiose (J) and sucrose (K) is colony size relative to growth on glucose. Strains (points) are colored according to whether they came from grapevine, oak-forest or oak-vineyard samples (source) and boxplots show median and interquartile range. Species were ordered with respect to mean normalized colony size.

